# Brain image-derived phenotypes yield insights into causal risk of psychiatric disorders using a Mendelian randomization study

**DOI:** 10.1101/2021.03.25.436910

**Authors:** Jing Guo, Ke Yu, Yan Guo, Shi Yao, Hao Wu, Kun Zhang, Yu Rong, Ming-Rui Guo, Shan-Shan Dong, Tie-Lin Yang

**Affiliations:** Key Laboratory of Biomedical Information Engineering of Ministry of Education, Biomedical Informatics & Genomics Center, School of Life Science and Technology, Xi’an Jiaotong University, Xi’an, Shaanxi, P. R. China, 710049

**Keywords:** Psychiatric disorders, IDPs, Causality, Mendelian Randomization

## Abstract

Brain imaging-derived phenotypes (IDPs) serve as underlying functional intermediates in psychiatric disorders, both of which are highly polygenic and heritable. Observational studies have elucidated the correlation between IDPs and psychiatric disorders, yet no systematic screening of IDPs to confer the causal liability in such disorders. We conducted a bidirectional two-sample Mendelian randomization framework to explore the causality of 1,901 IDPs on 10 psychiatric disorders and established the BrainMR database (http://brainmr.online/idp2psy/Index.php). We identified 17 causalities in forward MR analyses and 14 causalities in reverse MR analyses, which all rigorously examined them through various sensitivity analyses. The top significant relationship in forward was a unit lower volume of right central lateral of the thalamic nuclei, which was causally associated with an increased anorexia nervosa risk with an estimated OR of 0.52 (95% CI 0.41-0.64, *P* = 3.32 × 10^−9^). In reverse, the top significant IDP to be affected was the area of superior segment of the circular sulcus of the insula in the right hemisphere, which was increased by the reason of SCZ risk with an estimated association effect size of 0.068 (95% CI 0.046-0.090, *P* = 1.12 × 10^−6^). Overall, our study provides unique insights into causal pathways of psychiatric disorders at the imaging levels.

## Introduction

Psychiatric disorders are a group of serious brain dysfunction illnesses in which individuals are disturbed emotionally, cognitively or behaviorally. Millions of people worldwide are reported to suffer from psychiatric disorders, which are the main cause of disability and social burdens, and are therefore considered serious public health problems^1,2^.

Psychiatric disorders can be characterized by a collection of physiological brain changes, including many heritable image-derived phenotypes (IDPs) that can be measured non-invasively using magnetic resonance imaging (MRI)^3-6^. IDPs could serve as intermediate phenotypes between genomics and psychiatric disorders^7^. A large number of observational studies have been performed to explore the relationship between IDPs and psychiatric disorders. For example, the smaller cortical thicknesses of the insula and rostral anterior cingulate cortex are regarded as cerebral markers for adolescent anxiety^8^. The fractional anisotropy (FA) values of left superior longitudinal fasciculus, left prefrontal cortex, and left parietal lobe in patients with major depressive disorder (MDD) are lower than those in controls^9^. The thicknesses of the temporal and parietal cortex are decreased in patients with schizophrenia^10^, whereas increased in patients with autism spectrum disorder (ASD)^11^. However, single brain IDP may not accurately reflect the state of the disease in most cases, multiple IDPs are needed as diagnostic reference. Moreover, findings from conventional observational studies could not account for confounding factors. It is critical to explore the causal relationships between IDPs and psychiatric disorders.

Mendelian randomization (MR) is a widely used epidemiological method that uses genetic variants as instrument to infer the causality between exposure factors and outcomes^12,13^. With the accomplishment of the genome-wide association study (GWAS) on IDPs^14^, it provides us with the opportunity to systematically explore the causal relationships between all the IDPs and psychiatric disorders using large-scale data. Therefore, in this study, we performed an optimized two-sample MR framework to estimate causal risk factors for psychiatric disorders at brain IDPs levels. First, we systematically screened the causal effects of a total of 1,901 IDPs as exposures (including 1451 structural MRI and 450 diffusion MRI, respectively), on 10 psychiatric disorders as outcomes in almost 40,000 UK Biobank participants. Second, we assessed the commonality and specificity of brain IDPs, which could explain their wide-ranging effects in psychopathology. Finally, we established a BrainMR database (http://brainmr.online/idp2psy/Index.php) to provide a useful resource to share our results. Our findings could offer a systemic view of the causal effects of IDPs and psychiatric disorders. The identified IDPs may be applied as potential targets for etiology diagnosis.

## Results

### Overview of causality framework between IDPs and psychiatric disorders

We used an optimized two-sample MR pipeline to assess the bidirectional causal associations between 1,901 IDPs and 10 psychiatric disorders by using the GWAS summary-level data. The detailed information of all the GWAS statistics is summarized in Supplementary Tables 1-2. The flowchart of our analyses is shown in Figure 1.

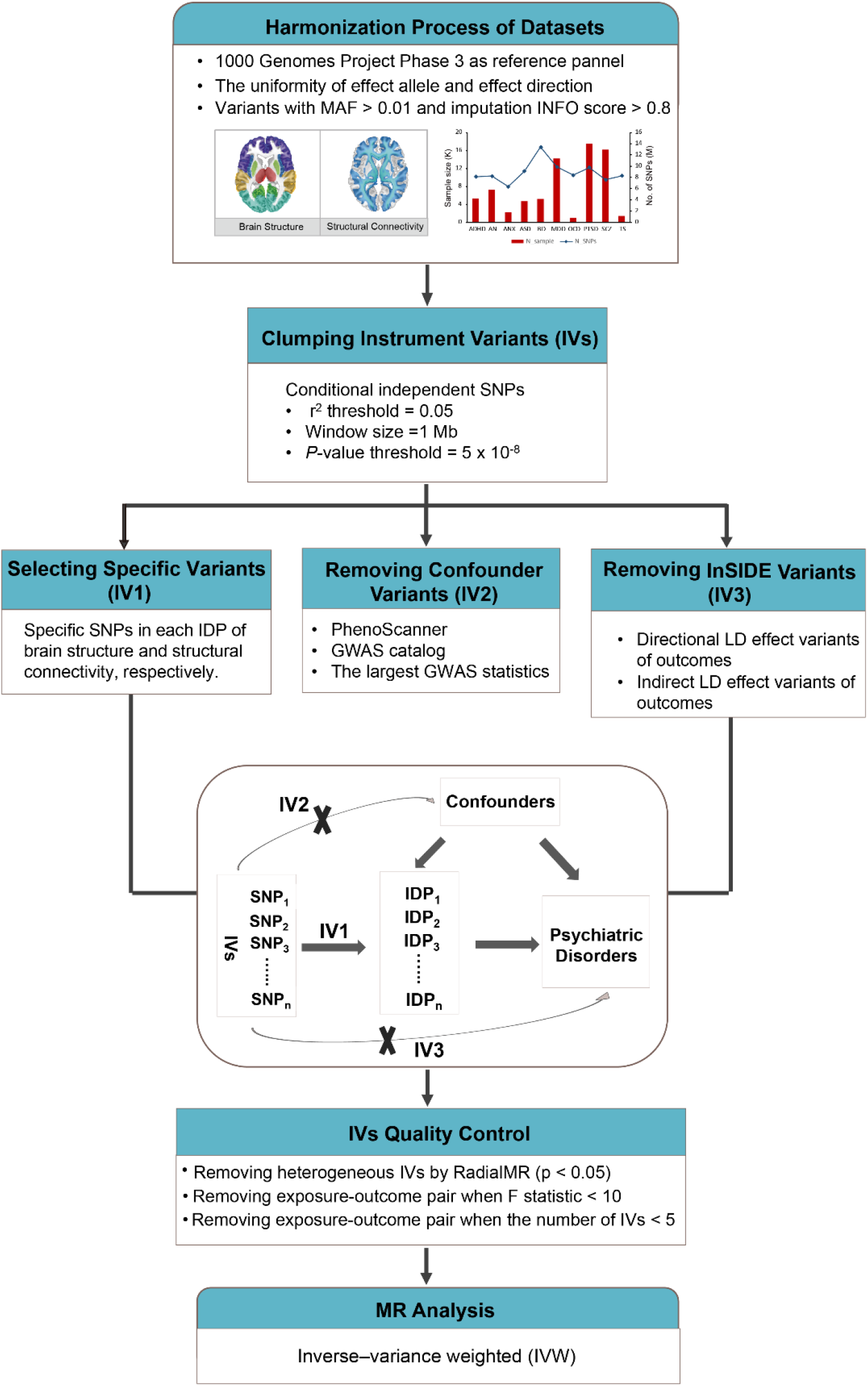

Severe standards were applied to select qualified instrument variants (IVs). We firstly clumped conditional independent SNPs for each IDP, but none of the 102 IDPs had any significant correlated common variants (*p* < 5 × 10^−8^). Then, we removed a total of 855 IVs in sum of 1044 IDPs, which were dependent of confounding effects, such as education, socioeconomic, smoking and drinking (Supplementary Table 3). We discarded 298 IVs that fell into the genome with direct or LD indirect influence on corresponding outcomes (292 IVs were associated with SCZ, 4 with AN and 2 with BD). We next conducted heterogeneity tests, excluding a total of 4,195 IVs as outliers from 5,720 exposure-outcome pairs. We finally obtained 10,158 exposure-outcome pairs, which meet the requirements of the strong instruments for a minimum number of five IVs and a valid F-statistic above the threshold of 10 (Supplementary Table 4). The final identified IVs of these exposure-outcome pairs were listed in Supplementary Table 5.

### Identification of the causal IDPs in psychiatric disorders

Since the majority of IDPs were genetically correlated with each other (Supplementary Table 6), we used the spectral decomposition method to provide a more appropriate estimate of the number of independent IDPs. Therefore, across 1047 independent IDPs, we selected a Bonferroni-adjusted *p*-value threshold of 4.78 × 10^−6^ for prioritizing MR results (number of exposure-outcome pairs = 10470). We exploited inverse-variance weighted (IVW) analysis as the primary method to estimate causal effects of each IDP on each of psychiatric disorders (full results shown in Supplementary Table 6). We ultimately identified 17 statistically significant causal relationships between 16 IDPs and 5 psychiatric disorders (Figure 2a). The numbers of causal IDPs from high to low involved in disorders were bipolar disorder (6), anorexia nervosa (4), SCZ (4), PTSD (2) and autism spectrum disorder (ASD) (1).

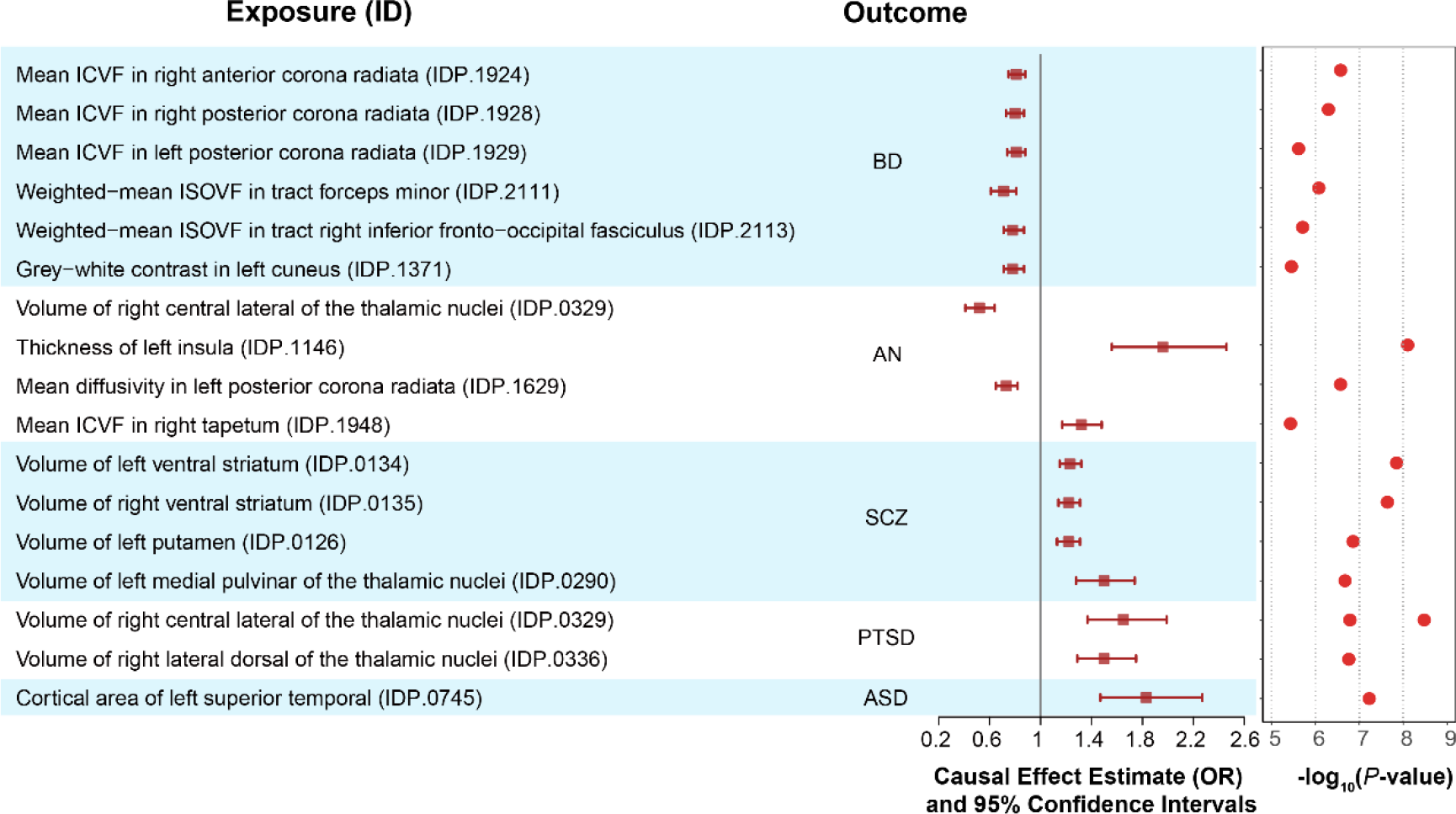

The specific results are set out as follows: Influential IDPs affected BD were all related to brain white matter microstructures with negative causalities, among them, the OR of estimation was ranged from 0.71 to 0.81. These causal IDPs including: three projection fibers of mean ICVF (intra-cellular volume fraction) in the right anterior (OR = 0.81, 95% CI 0.75-0.88, *P* = 2.66 × 10^−7^) and the bilateral posterior (right: OR = 0.80, 95% CI 0.73-0.87, *P* = 5.04 × 10^−7^; left: OR = 0.81, 95% CI 0.74-0.88, *P* = 2.41 × 10^−6^) corona radiata, which involved in the limbic-thalamo-cortical circuits; one commissural fibers of weighted-mean ISOVF (isotropic or free water volume fraction) in the forceps minor tracts (OR = 0.71, 95% CI 0.61-0.81, *P* = 8.31 × 10^−7^), which is an important connection between the prefrontal lobes of the two hemispheres of the brain; one association fibers of ISOVF in the right inferior fronto-occipital fasciculus (OR = 0.78, 95% CI 0.71-0.87, *P* = 1.95 × 10^−6^), which connected the frontal lobe to posterior basal temporal and inferior occipital gyrus; and one grey-white contrast indicated the alterations of myelin sheath in cuneus of the left hemisphere (OR = 0.78, 95% CI 0.71-0.87, *P* = 3.51 × 10^−6^). Anorexia nervosa was detected with 4 causal IDPs, of which 2 positive causalities and 2 negative causalities. The top significant relationship was a unit lower volume of right central lateral of the thalamic nuclei, which was causally associated with an increased anorexia nervosa risk with an estimated OR of 0.52 (95% CI 0.41-0.64, *P* = 3.32 × 10^−9^). There was a projection fiber in the limbic-thalamo-cortical circuits that we mentioned above, which was negatively correlated with anorexia nervosa (OR = 0.73, 95% CI 0.65-0.82, *P* = 2.67 × 10^−7^), that is, the mean diffusivity in the left posterior corona radiata. The insula, which is located deep in the lateral sulcus, was identified its higher thickness in left hemisphere associated with an increased anorexia nervosa risk (OR = 1.96, 95% CI 1.56-2.46, *P* = 7.90 × 10^−9^). A commissural fiber of mean ICVF in right tapetum was positively causal associated with anorexia nervosa (OR = 1.32, 95% CI 1.17-1.48, *P* = 3.68 × 10^−6^), which connected the inferior temporal lobes on both sides of brain hemisphere via the splenium of the corpus callosum. For SCZ, we identified the volumes of four brain anatomical structure were as risk factors, namely the bilateral ventral striatum (right: OR = 1.22, 95% CI 1.14-1.31, *P* = 2.29 × 10^−8^; left: OR = 1.23, 95% CI 1.15-1.32, *P* = 1.41 × 10^−8^), left putamen (OR = 1.22, 95% CI 1.13-1.31, *P* = 1.39 × 10^−7^), and medial pulvinar of the thalamic nuclei (OR = 1.50, 95% CI 1.28-1.74, *P* = 2.11 × 10^−7^), which were participated in cortico-basal ganglia-thalamocortical-cortical circuits. The effect of the right central lateral of the thalamic nuclei on PTSD was opposite to anorexia nervosa, which increased PTSD risk with a unit higher volume on an effect of the estimated OR of 1.65 (95% CI 1.37-1.99, *P* = 1.63 × 10^−7^). The volume of another region of thalamic nuclei in the right lateral dorsal was also increased the PTSD risk with an estimated OR of 1.50 (95% CI 1.29-1.75, *P* = 1.73 × 10^−7^). Only one significant causal IDP was identified for ASD, the cortical area of left superior temporal, which was a risk factor when increased with an estimate OR of 1.83 (95% CI 1.47-2.27, *P* = 5.91 × 10^−8^).

We performed sensitivity analyses to validate the causal effects of IDPs on the main MR results. First, no horizontal pleiotropy was found by using the global test of MR-PRESSO (Supplementary Figure 1a). Second, the Egger intercepts were almost close to zero in all analyses except one relationship, between the volume of the left ventral striatum and SCZ (Intercept = -0.013, *p*-value = 0.049), which was probably disputed with directional pleiotropy (Supplementary Figure 1b). Third, we conducted leave-one-out cross-validation, and found no outliers (Supplementary Figure 2). Additionally, we used another three MR approaches, including MR-Egger regression, weighted median-based MR and mode-based MR to examine our results of IVW method. We identified that the main MR results were successfully confirmed by at least one of the three methods with the consistent directions (normal *p*-value < 0.05) (Supplementary Figure 3).

### Reverse MR between genetically determined psychiatric disorders and IDPs

A total of 7,318 exposure-outcome pairs available for MR analyses after performing our predefined IVs selection criteria. The exposures were mainly clustering in BD, AN, SCZ and ADHD, and their corresponding IVs were shown in the Supplementary Table 8. Since we still set the same significance threshold based on the effective number of independent IDPs, 14 statistically significant causal relationships affected by SCZ were identified (Figure 3, and full results shown in Supplementary Table 9). Among them, the estimated effect was ranged from -0.067 to 0.068, and the *p*-value of significant causality was ranged from 1.12× 10^−9^ to 4.32 × 10^−6^. According to the MR results, we could audacious speculate the pathogenesis of SCZ might occur the microstructure alterations in six anatomical parts of brain, including: insula, frontal lobe, temporal lobe, occipital lobe, limbic system and projection fibers of white matter. For example, the top significant IDP to be affected was the area of superior segment of the circular sulcus of the insula in the right hemisphere, which was increased by the reason of SCZ risk with an estimated association effect size of 0.068 (95% CI 0.046-0.090, *P* = 1.12 × 10^−6^). Taken together, the development and pathology of schizophrenia probably be different pathways in the light of the interpretation between forward and reverse MR results.

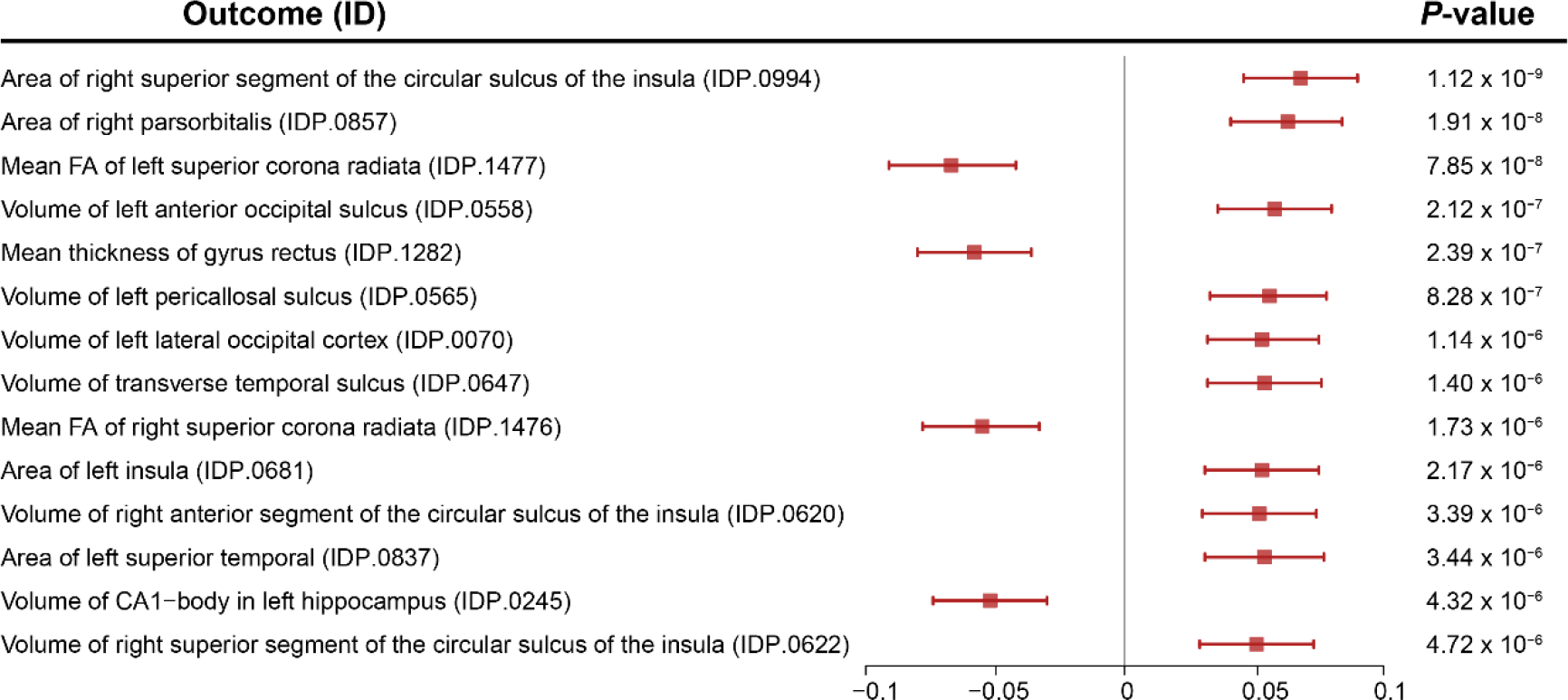

Sensitivity analyses suggested the robust evidence to support our reverse MR results without horizontal pleiotropy (Supplementary Figure 4a) or outliers (Supplementary Figure 5). However, 5 IDPs did not hold in the MR-Egger regression analyses, whose intercepts were significantly deviated from zero because of directional pleiotropy (Supplementary Figure 4b). Confidently, the primary IVW results were accordant with those of other MR methods, especially the weighted median-based MR ((Supplementary Figure 6).

All of our results of causality between brain IDPs and psychiatric disorders were integrated into an open online BrainMR database (http://brainmr.online/idp2psy/Index.php) for public access. All the raw data in the BrianMR database have been uniformly stored, so as to facilitate rapid and systematic queries.

## Discussion

In this study, we performed two-sample MR analyses to systematically investigate the causal effects of 1,901 IDPs on 10 psychiatric disorders, with large GWAS statistics from UK biobank and PGC database. Most of the previous studies were based on imaging data from case-control groups, which was limited by the sample size, high economic cost, and especially lack of causal reasoning. In addition, the types of imaging data reported in previous studies were relatively limited. We identified a total of 17 pairs of causal IDPs on psychiatric disorders, which could be used to prioritize promising biomarkers for predicting the risk of psychiatric disorders.

Influential IDPs affected BD with the largest number, all of which were involved in brain white matter microstructure. Patients with BD are characterized by recurrent mania and depression symptoms during the inter-episodic period. Numerous researches have commonly demonstrated that brain white matter activity make an important impact on BD pathology^15,16^. In our causation findings, negative causal factors of BD were identified in grey-white contrast in cuneus of left hemisphere, mean ICVF (intra-cellular volume fraction) in right anterior corona radiata and bilateral posterior corona radiata, and weighted-mean ISOVF (isotropic or free water volume fraction) in tract forceps minor and right inferior fronto-occipital fasciculus. Previous researches have shown the supported information in line with the most results of our present study. For example, the inferior fronto-occipital fasciculus is one of the association fibers connecting from the frontal lobe to posterior basal temporal and inferior occipital gyrus#1, which is demonstrated that associated with reduced FA in BD subjects compared to health control populations^17^. Studies on the function of the inferior fronto-occipital fasciculus also implicate the interaction between visual and emotional functions^18^ to regulate the facial emotional recognition and responding^19^. It has been widely demonstrated that frontal lobe is a crucial area of BD in the field of brain imaging research. Among the regions connected, the frontopolar cortex has been shown to be important for cognitive behaviors in human and non-human primates. The deficient modulation in frontal regions, especially in white matter integrity, have been demonstrated to influence the emotional dysregulation in BD^20,21^. The observed causal inducer of BD, forceps minor, which is one of the commissural fibers that connects the prefrontal cortex of interhemispheric with crossing the genu of the corpus callosum. At present, there is less study to forces minor abnormality and BD, but forces minor is an important connection between the prefrontal lobes of the two hemispheres of the brain. Previous research has shown that the forceps minor is implicated in cognitive dysfunctions, and the FA values in the forceps minor can be used for prediction of cognitive impairments in patients^22^, and higher diffusion properties of the forceps minor were important for learning and attention skills^23^.

Integrating all the causal inference risk factors of BD, we can find that these risk IDPs are mainly concentrated in the white matter fiber tracts from frontal lobe to corpus callosum, occipital lobe and temporal lobe, thus transmitting information between these regions. These areas have been proved to be related to cognition, attention-control, emotion and speech-language. Although the sample size of most of previous studies is small, they are consistent with direct or indirect evidence to support our results.

Our findings suggest that IDP has a specific effect on the psychiatric disorder, which is unlike the similar polygenic overlap shared among these diseases. A causal IDP might be related to one disease, which indicated an important risk factor leading to the occurrence of this disease. Moreover, there are multiple IDPs causally associated with the same psychiatric disorder. We could integrate multiple IDPs as risk evaluation indicators for pathological occurrence, which could provide theoretical support for disease prediction and diagnosis.

## Online Methods

### Data sets

#### Exposure data -brain IDPs

1,901 IDPs were selected as exposures, which were divided into two categories including brain structure (structure MRI) and brain structural connectivity (diffusion MRI) and each was in up to 40,000 European ancestry subjects. Brain structure included 1451 anatomical subtypes such as volume, superficial area and subcortical thickness. Brain structural connectivity included 450 subtypes of long-range structural connectivity and local microstructure with a series of orientation signals by diffusion-weighted imaging (dMRI). The detailed information is provided by UK Biobank online refs (https://biobank.ctsu.ox.ac.uk/crystal/crystal/docs/brain_mri.pdf). Briefly, original IDPs of brain structure were T1 or T2 weighted structural images^24^, and different algorithms or methods were performed to generate new image features. Brain structural connectivity IDPs were initially screened by dMRI that was a technique to detect water molecules diffusion with a range of orientations within local region^25^. All GWAS summary statistics of IDPs were collected from the Oxford BIG web browser (http://big.stats.ox.ac.uk/)3.

#### Outcome data -psychiatric disorders

We collected ten psychiatric disorders with publicly available GWAS summary statistics as outcomes, namely ADHD^26^, anorexia nervosa^27^, ASD^28^, bipolar disorder^29^, MDD^30^, OCD^31^, PSTD^32^, schizophrenia^33^, Tourette syndrome^34^ and anxiety^35^. The vast majority of these data were derived from Psychiatric Genomics Consortium (PGC)^36^. Individuals used in outcomes were independent of exposures, the detailed information is summarized in Supplementary Table 2.

#### Harmonized process of datasets

To guarantee the precise causal inference, we conducted harmonization process of original GWAS statistics, which could check whether the SNP alleles were coded from the same strand of DNA^37^. We took genomic reference of 1000G for allele matching. The minor alleles were defined as the effect alleles, which were matched to reference genomes to ensure uniformity of effect direction. Moreover, we checked for palindromic SNPs whose alleles correspond to nucleotides that pairs in forward and reverse coding, such as A/T and G/C. We discarded SNPs if their minor allele frequencies (MAFs) were close to 0.50 (e.g., 0.48 -0.52), which might introduce ambiguity into the conformity of the effect allele between the exposure and outcome datasets^38^. About 0.20% -0.47% palindromic SNPs were deleted (Supplementary Table 9).

### MR Analysis

#### Selection of instrument variants

Firstly, conditional independent SNPs of each IDP were clumped by using PLINK with a stepwise model-selection procedure^39^. We set linkage disequilibrium (LD) pruning r2 threshold = 0.05 in UK Biobank reference, window size = 1 Mb and p-value threshold = 5 × 10^−8^. Then, IVs should be associated with target IDP exclusively and not associated with other IDPs in one category (brain structure or connection) to precluded pleiotropic variants. IVs should satisfy three assumptions of conventional MR principle, that is, it must be statistically significant associated with exposure, but not associated with any confounder, and not associated with outcome. We also excluded the variants whose LD was directly or indirectly related to outcome variants with the option (p-value < 5 × 10-8 and r2 > 0.8). We set p-value < 5 × 10^−8^ in the step of calculating conditional independent SNPs as IVs (Supplementary Figure 5a-b).

#### Removing confounders

In order to minimize pleiotropy, we removed the multiple IVs associated with confounders. We used PhenoScanner GWAS database (http://www.phenoscanner.medschl.cam.ac.uk/) and NHGRI-EBI GWAS catalog (https://www.ebi.ac.uk/gwas/) to assess any previous associations with confounders. We also excluded confounder IVs directly associated with alcohol intake44 and smoking45, because these factors could affect brain imaging phenotypes.

#### Quality control of instrument variants

In order to improve the accuracy and robustness of genetic instruments, we proposed heterogeneity test to detect outliers and adjusting them. We used Cochran’s Q test for Inverse variance weighted (IVW) model fit in the condition of horizontal pleiotropic influences and exploited Rucker’s Q’ test for MR-Egger fit by reason of directional pleiotropic influences47,48. We implemented the RadialMR version 0.4 R package to calculate the modified Q and Q’ test^40^, and discarded the statistically significant outliers at the nominal significance level of 0.05. Additionally, we performed fist-stage F-statistics to measure the power of the final genetic instruments, of which lower F statistics correspond to greater bias. We set a conventional threshold of 10 that meaning a qualified IVs set and weak instrument bias is unlikely to occur in MR analysis. We also discarded the exposure-outcome MR test when the number of IVs less than five.

#### Bi-directional MR analyses

We performed two-sample MR analyses to explore the causal relationships between brain IDPs and psychiatric disorders with the valid IVs. Inverse variance weighted (IVW) regression was conducted as the primary causal infere^41^. Since IVW might be biased if any of the SNPs exhibit horizontal pleiotropy, we applied other MR analysis as complementary methods to strengthen the confidence. MR-Egger method estimates the causal effect by slope coefficient of Egger regression, which can provide a more robust estimate even when all genetic variants are invalid IVs^42^. Weighted median-based estimator could offer protection against even up to 50% invalid IVs^42^. Mode-based method provides a consistent estimator when relaxed IV assumptions with less bias and lower type-I error rates^43^. All these methods were implemented in the TwoSampleMR version 0.4.10 R package^44^.

#### Sensitivity Analyses

For outcomes with significant MR analysis results, we conducted sensitivity analyses. First, we executed a leave-one-out cross-validation analysis to check whether causal association was driven by a single SNP. Second, we conducted MR-PRESSO (Mendelian randomization pleiotropy residual sum and outlier) test to detect whether there was any horizontal pleiotropic outlier in our MR study^45^. P value less than 0.05 was regarded as an outlier in the statistical significance test. Third, we performed MR-Egger regression to check for potential bias from directional pleiotropy. The intercept in the Egger regression represents the average pleiotropic effect of all genetic variants, and is interpreted as evidence of directed pleiotropy when the value differs from zero. Fourth, to guard against the small possibility that our results were driven by reverse causality, we performed IVW analyses in reverse direction to examine the possible reverse causation bias from psychiatric disorders to IDPs. In this study, we conducted MR in accordance with Strengthening the Reporting of Observational Studies in Epidemiology (STROBE) and STROBE-MR guidelines^46^.

## Data availability

The full GWAS summary-level data of brain IDPs were collected from Oxford BIG web browser (version 2.0) through http://big.stats.ox.ac.uk/download_page. GWAS summary statistics of ADHD, AN, ANX, ASD, BD, MDD, OCD, PTSD and TS from PGC are available on the Psychiatric Genomics Consortium’s downloads page (https://www.med.unc.edu/pgc/download-results). SCZ was collected from CLOZUK and PGC2 meta-analysis summary statistics through https://walters.psycm.cf.ac.uk. Summary statistics of the rest of the analyzed traits for condounders removing analyses are available at GRASP (https://grasp.nhlbi.nih.gov/FullResults.aspx), or Gene ATLAS (http://geneatlas.roslin.ed.ac.uk/downloads), or CNCR (https://ctg.cncr.nl/software/summary_statistics), or CCACE (https://www.ccace.ed.ac.uk/node/335), or epiGAD (http://www.epigad.org/gwas_ilae2014), or CDKP (https://cd.hugeamp.org/downloads.html), or http://cnsgenomics.com, or GIANT (http://portals.broadinstitute.org/collaboration/giant/index.php/GIANT_consortium_data_files).

## Code availability

LD-based result clumping was performed using PLINK 1.9 (https://www.cog-genomics.org/plink/1.9/). Genetic correlation analysis was performed using LDSC (https://github.com/bulik/ldsc). R version 3.3.2 (https://www.r-project.org/) was used for statistical analysis. The TwoSampleMR v0.4.10 R package (https://mrcieu.github.io/TwoSampleMR/), RadialMR v0.4 (https://github.com/WSpiller/RadialMR/) and MR-PRESSO (https://github.com/rondolab/MR-PRESSO) were used for MR analyses.

PhenoScanner GWAS database (http://www.phenoscanner.medschl.cam.ac.uk) and NHGRI-EBI GWAS catalog (https://www.ebi.ac.uk/gwas/) were used to remove confounding factors.

## Acknowledgements

This work was supported by grants from the National Natural Science Foundation of China (31871264, 31970569), the Natural Science Basic Research Program Shaanxi Province (2019JM-119) and the Fundamental Research Funds for the Central Universities. We would like to thank the UK Biobank and Cross-Disorder Group of the Psychiatric Genomics Consortium (PGC).

## Author contributions

T.L.Y. conceived and supervised this project. J.G. conducted the computational work and wrote the manuscript. K.Y built the website of database. Y.G. revised the manuscript. S.Y and other authors offered some advices and assistances.

